# mir-71 mediates age-dependent opposing contributions of the stress activated kinase KGB-1 in *Caenorhabditis elegans*

**DOI:** 10.1101/835355

**Authors:** Cyrus Ruediger, Michael Shapira

## Abstract

Studying the evolutionary processes that shaped aging offers a path for understanding the causes of aging. The Antagonistic Pleiotropy theory for the evolution of aging proposes that the inverse correlation between natural selection strength and aging allows positive selection of gene variants with early-life beneficial contributions to fitness in spite of detrimental late-life consequences. However, mechanistic understanding of how this principle manifests in aging is still lacking. We previously identified antagonistic pleiotropy in the function of the *Caenorhabditis elegans* JNK homolog KGB-1, which provided stress protection in developing larvae, but sensitized adults to stress and shortened their lifespan. To a large extent, KGB-1’s contributions depended on age-dependent and opposing regulation of the stress transcription factor DAF-16, but the underlying mechanisms remained unknown. Here we describe a role for the microRNA mir-71 in mediating effects of KGB-1 on DAF-16 and on downstream phenotypes. Fluorescent imaging along with genetic and survival analyses revealed age-dependent regulation of *mir-71* expression by KGB-1 – upregulation in larvae, but downregulation in adults, and showed that *mir-71* was required both for late-life effects of KGB-1 (infection sensitivity and shortened lifespan), as well as for early life resistance to cadmium. While *mir-71* disruption did not compromise development under protein folding stress (known to depend on KGB-1), disruption of the argonaute gene *alg-1*, a central component of the microRNA machinery, did. These results suggest that microRNAs play a role in mediating age-dependent antagonistic contributions of KGB-1 to survival, with mir-71 playing a central role and additional microRNAs contributing redundantly.

## INTRODUCTION

Pleiotropic effects manifesting at different ages are the basis of the Antagonistic Pleiotropy theory for the evolution of aging. This theory proposes that since the strength of natural selection declines with age, gene variants with late-life deleterious effects can still be positively-selected if they have early-life beneficial effects (Williams 1957). This theory explains aging as the result of gene variants that promote early life processes, such as development and reproduction, but delimit lifespan. Examples of antagonistic pleiotropy have been described (and debated), including the TOR pathway, which regulates protein synthesis with contrasting impacts on growth and development versus lifespan, and p53 which prevents cancerous cell proliferation, but also promotes cell senescence (Rodríguez *et al.* 2017; Long and Zhang 2019; Ungewitter and Scrable 2009; Kapahi *et al.* 2010). However, mechanistic understanding of when and how a good contribution becomes bad is still missing.

We previously identified age-dependent contributions for the *Caenorhabditis elegans* c-jun N-terminal kinase homolog KGB-1, which demonstrated characteristics of antagonistic pleiotropy (Twumasi-Boateng *et al.* 2012). KGB-1 is expressed throughout development and adulthood in various tissues, including the gut, epidermis, muscle, neurons, and the gonad (Liu *et al.* 2018). Early work demonstrated its importance for reproduction (Orsborn *et al.* 2007). Work utilizing a hypomorphic allele further showed that KGB-1 also provided protection from heavy metals and protein misfolding stress (ER stress), enabling development under adverse environmental conditions (Mizuno *et al.* 2004, 2008). More recently, we found that these protective contributions of KGB-1 were limited to developing larvae, whereas its activation in adults (following knock-down of a negative regulator, the phosphatase VHP-1) increased sensitivity to the same stresses, as well as to infection, and shortened lifespan. While VHP-1 is also a negative regulator of P38 MAPK signaling, the described effects on stress resistance were independent of the p38 pathway (Twumasi-Boateng *et al.* 2012). To a large extent, age-dependent contributions of KGB-1 depended on the stress-protective and longevity-associated transcription factor DAF-16 (Kenyon *et al.* 1993; Larsen *et al.* 1995; Ogg *et al.* 1997). In larvae, KGB-1 activation promoted DAF-16 nuclear localization and transcriptional output, while the same activation in young adults attenuated nuclear localization and output (Twumasi-Boateng *et al.* 2012). A second transcription factor, FOS-1, was found to be important for gene expression downstream of KGB-1, but its contributions were age-invariant (Zhang *et al.* 2017).

How KGB-1 modulated DAF-16 was not clear. Previous results indicated this could be achieved cell nonautonomously, with neuronal KGB-1 modulating intestinal DAF-16 nuclear localization (Liu *et al.* 2018), but to date, no molecular mechanism was identified that mediated these effects. Here, we show that the microRNA mir-71, and potentially additional microRNAs, are regulated by KGB-1 in an age-dependent manner and are required to mediate effects on DAF-16, as well as for a subset of KGB-1 age-dependent contributions to stress resistance and lifespan.

## MATERIALS AND METHODS

### Strains

Strains used included *kgb-1(km21), alg-1(tm492), alg-2(ok304)*, TJ356 (*zIs356 [daf-16p::daf-16a/b::GFP + rol-6(su1006)]*), *drsh-1(ok369)/hT2 [bli-4(e937) let-?(q782) qIs48]), pash-1(mj100);mjEx331[eft-3p::pash-1::GFP::unc-54 3’UTR;myo-2p::mCherry::unc-54 3’UTR], mir-71(n4115), daf-12(rh61rh411), kri-1(ok1251)* and the N2 wild-type strain. All were obtained from the *Caenorhabditis* Genetics Center (Minneapolis, MN). *pash-1(mj100)* and *drsh-1(ok369)* mutants were isolated from genetically rescued or balanced strains, respectively, by picking non-fluorescent animals. Strains containing combinations of the aforementioned mutations and transgenes were generated by mating and verified by PCR and sequencing. *unc-119(ed3);maIs352[mir-71p::GFP;unc-119(+)]* worms were gratefully received from Dr. Zachary Pincus, Washington University.

Bacterial strains included *Escherichia coli* OP50-1, and *Pseudomonas aeruginosa* PA14 (Shapira and Tan 2008).

### Imaging

Worms were picked onto unseeded agarose plates, paralyzed with 25 mM levamisole (Sigma, St. Louis, MO), and imaged using a Leica MZ16F fluorescent stereoscope (Leica, Wetzlar, Germany) equipped with a MicroPublisher 5.0 RTV (QImaging, BC, Canada). Images were processed and quantified using ImageJ (National Institutes of Health, Bethesda, MD).

### RNA interference-mediated knockdown

Knockdown by feeding was performed using standard protocols, with clones from the Ahringer RNAi library (Kamath et al. 2003), except for the vhp-1 clone, which was from the Open Biosystems Library (Reboul et al. 2003). Bacteria harboring an empty RNAi vector (EV) served as control. Larval knockdown was achieved by exposing worms from the egg stage to larval L3 or L4, as described; knockdown in adults was carried out from L4 to day 2 of adulthood, unless otherwise mentioned. All experiments were performed at 20°C, unless otherwise specified.

cdc-25.1 RNAi was used to disrupt germline proliferation, as described elsewhere (Shapira and Tan 2008), sterilizing worms and promoting DAF-16 nuclear localization downstream of gonad signaling. Young adult parents were fed cdc-25.1 RNAi for 8 hours and their progeny further grown on cdc-25.1 RNAi until the L4 larval stage, and then transferred to plates containing a 1:1 mixture of cdc-25.1 RNAi with vhp-1 RNAi or empty vector clones.

### Acute Cadmium Toxicity Resistance Assays

Worms were fed EV or vhp-1 RNAi from hatching until the L3 stage, then transferred to K media plates [1.55 g NaCl, 1.19 g KCl, 8.5 g Bacto™ Agar, H_2_O to 1 liter; sterilized by autoclaving] with 10 mM CdCl_2_ and seeded with OP50-1. Survival was scored after 11 hours, at which point the majority of EV fed worms were dead.

### Tunicamycin development assays

Eggs were transferred to NGM plates containing 1μg/ml tunicamycin. After 3 days at 20°C, worms were staged and counted, and the percentages at different stages, as well as dead worms, calculated.

### Survival assays

Survival experiments were carried out at 20°C, unless otherwise specified. Worms were exposed to RNAi from L4 to day 2 of adulthood, and then transferred to NGM plates with 100 μg/ml kanamycin seeded with kanamycin-killed OP50-1, and survival scored every 2 days. Statistical evaluation of differences between survival curves was performed using Kaplan–Meier analysis, followed by a log-rank test.

### Quantitative RT-PCR

Total RNA was extracted from 100 to 500 animals using TRIzol (Invitrogen). RNA was treated with DNase to remove genomic DNA contamination (QIAGEN, Hilden, Germany), and cDNA was synthesized using iScript™ (Bio-Rad). SYBR Green quantitative (q)RT-PCR was performed using the SsoAdvanced Universal SYBR Green Supermix (Bio-Rad) on a StepOnePlus system (Applied Biosystems, Foster City, CA). Gene-specific threshold cycle (Ct) values were normalized by subtracting the respective values for measurements of actin gene expression. Statistical significance was calculated with a t-test based on actin-normalized Ct values. Primers, whose sequences are listed below, were used with an annealing temperature of 60°.

mtl-1 forward: GAGGCCAGTGAGAAAAAATGCT

mtl-1 reverse: GCTCTGCACAATGACAGTTTGC

actin forward: TCGGTATGGGACAGAAGGAC

actin reverse: CATCCCAGTTGGTGACGATA

mir-71 forward: -CGACGGCGAAAAACAGAATA

mir-71 reverse: -GTCTGCTCTGAACGATGAAAG

### Data availability

Strains used in this study are available upon request. The authors affirm that all data necessary for confirming the conclusions of the article are present within the article, figures, and tables.

## RESULTS

### *mir-71* is required for detrimental effects of KGB-1 activation in adults

Previous results have shown that KGB-1 activation following *vhp-1* knock-down in adults decreased infection resistance and lifespan (Twumasi-Boateng 2012). This was shown to be due to attenuation of DAF-16 nuclear localization in adults, particularly apparent following germline disruption where *vhp-1* knock-down abolishes an otherwise prominent DAF-16 nuclear localization (Liu 2018). Effects of KGB-1 activation on survival of germline-disrupted animals are reproducible, but their extent variable, likely due to variable efficiency of RNAi-mediated gene knock-down. In some cases, vhp-1 knock-down suppressed the lifespan-extension effect of germline disruption to the extent that sterile and fertile worms had similar lifespans (Fig. 1A). This observation suggested that at least one mechanism through which KGB-1 regulated DAF-16 involved pathways activated by germline disruption. Candidates, known to contribute to lifespan extension following germline disruption and putatively affected by KGB-1 activation included the dafachronic acid binding nuclear hormone receptor *nhr-12* and *kri-1*, both previously shown to promote DAF-16 nuclear localization in germline ablated animals (Berman and Kenyon 2006; Gerisch *et al.* 2007), and *mir-71*, a microRNA shown to promote DAF-16 nuclear localization and extend lifespan putatively by inhibiting insulin signaling (de Lencastre *et al.* 2010; Boulias and Horvitz 2012). Of these, only *mir-71* disruption reproducibly prevented the detrimental effects of KGB-1 activation in germline-disrupted adults (Fig. 1B, Table S1). Disruption of *mir-71* largely prevented detrimental effects of *vhp-1* knock-down also in fertile animals, but not as reproducibly (Table S2).

**Figure 1.**
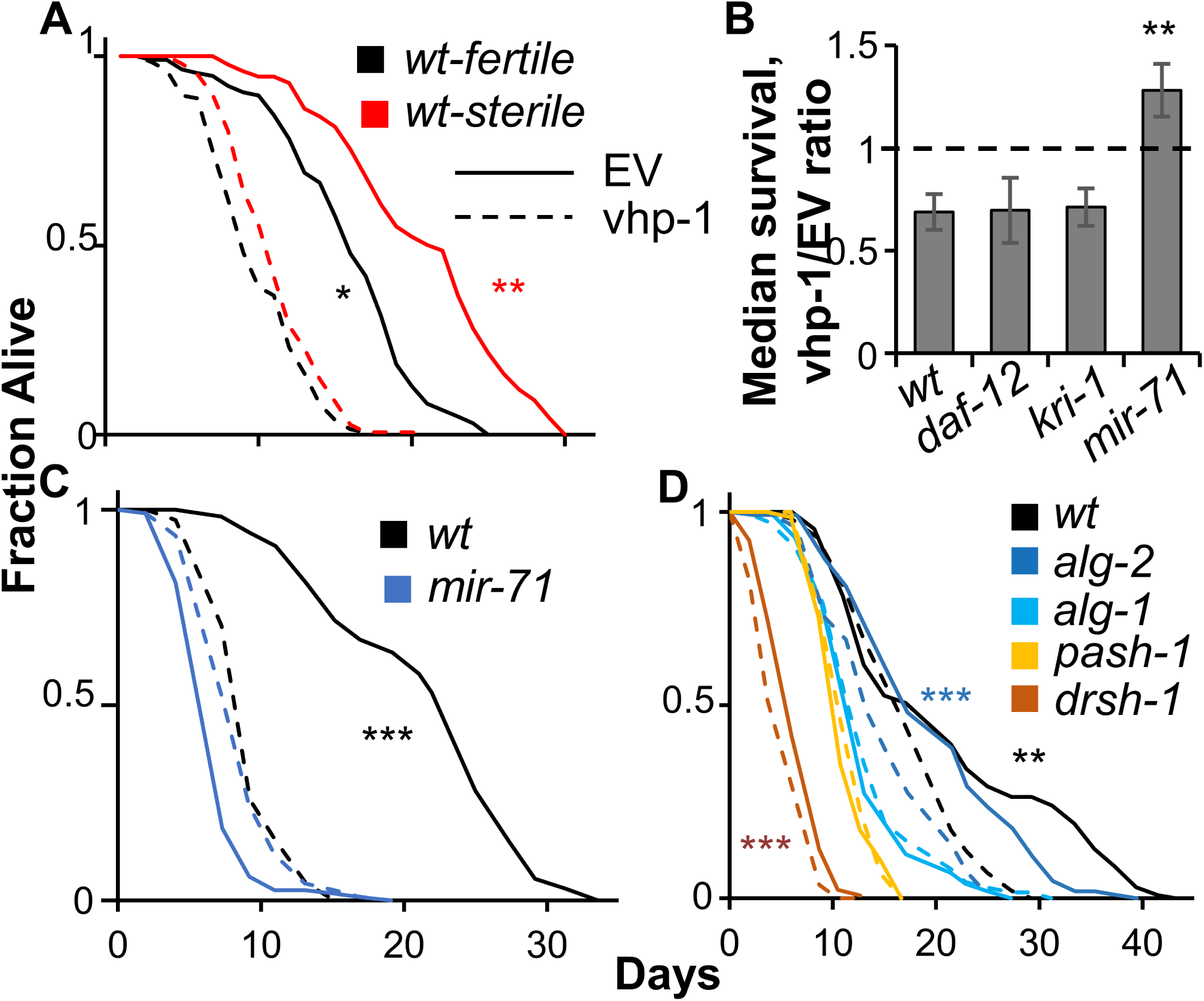
Detrimental effects of KGB-1 depend on mir-71. **(A)** Lifespan trajectories for wildtype animals, either fertile or sterilized by cdc-25.1 RNAi, exposed for two days starting at L4 to control (EV, empty vector), or vhp-1 RNAi (N=108-137 worms per group; *, p<0.05, **, p<0.01). **(B)** Ratios of median survival time of animals (of the designated strains, all cdc-25.1 RNAi-sterilized) exposed to vhp-1 RNAi during adulthood over the respective median of those exposed to control RNAi (EV). Included in median calculations for each strain are two *P. aeruginosa* infection survival assays, as well as one lifespan assay for wildtype and *mir-71* worms (see Table S1 for individual assays). **, p<0.01 compared to wildtype. (C) Lifespan assays for sterile wildtype and mir-71 animals, of a separate experiment than those included in B (N=81-118 worms per group, ***, p<0.001); shown is One representative experiment of three. (D) Lifespan assays of fertile microRNA processing mutants treated with control or vhp-1 RNAi as described above. Asterisks, as above (N=48-135 per group).

Argonaute-like proteins are required for microRNA processing and binding, and the two main *C. elegans* homologs, *alg-1* and *alg-2* were reported to have opposite effects on lifespan, with *alg-1* extending it and *alg-2* restricting it, both through interactions with IIS (Aalto 2018). We found that *alg-1* was required for the detrimental effects of KGB-1 activation, while *alg-2* was not (Fig. 1D). Furthermore, disruption of the *pash-1*, encoding a nuclear RNAase III co-factor necessary for microRNA processing (but not siRNA/RNAi), abolished the detrimental effects of KGB-1 activation. In contrast, animals with disruption of the RNAase III gene itself, *drsh-1*, which are short-lived to begin with, still showed detrimental effects following *vhp-1* knock-down.

### KGB-1 activation modulates mir-71 gene expression in an age-dependent manner

To examine the relationship between KGB-1 activation and mir-71 expression we employed transgenic worms expressing GFP from the *mir-71* promoter (Martinez *et al.* 2008). KGB-1 activation following *vhp-1* knock-down in adults resulted in a significant decrease in expression from the *mir-71* promoter, particularly in the intestine (Fig. 2). This was observed as early as two days from the beginning of RNAi exposure, but was more prominent following longer exposure, until day five of adulthood, when *mir-71p*-driven GFP expression is maximal (Pincus *et al.* 2011). Furthermore, suppression of *mir-71* expression was prominent in cdc-25.1(RNAi)-sterilized worms (Fig. 2), but was also apparent in fertile animals (not shown). In all cases, *kgb-1* disruption dramatically reduced the effects of *vhp-1* knock-down, supporting the notion that suppression of *mir-71* expression was KGB-1 dependent.

**Figure 2.**
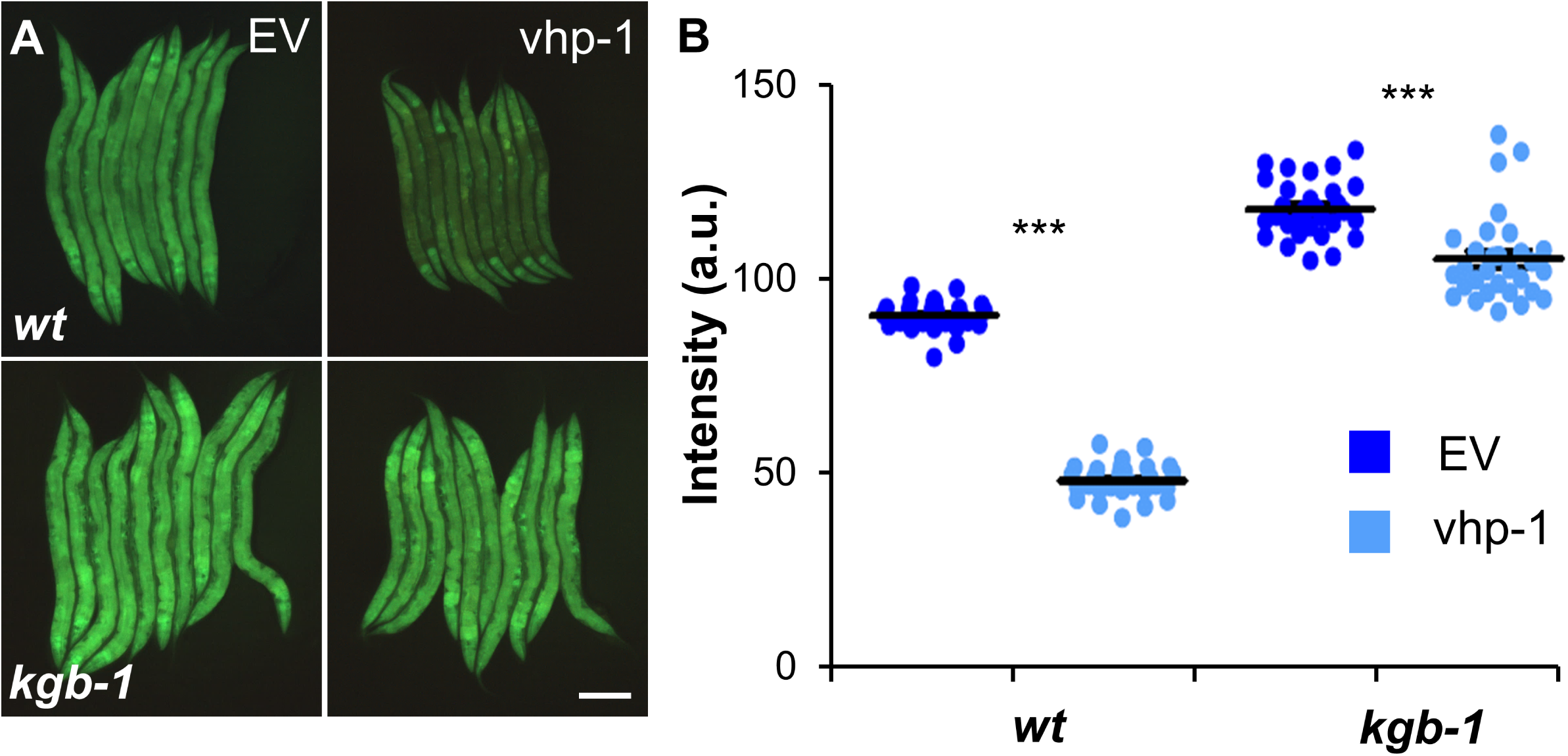
KGB-1 activation in adults suppresses *mir-71* expression. **(A)** Representative images of cdc-25.1(RNAi)-sterilized transgenic animals expressing GFP from the *mir-71* promoter, in wildtype or *kgb-1(km21)* background, and following a 5-day exposure, beginning at L4, to control (EV) or vhp-1 RNAi. Scale bar, 200 mm (B) Quantification of signal intensity from worms as in A. Lines mark averages ± SE; individual measurements shown in dots, 29-30 worms per group; *** p<0.001, t-test.

Previous results demonstrated that KGB-1 contributed to downstream gene expression and phenotypes mostly through cell non-autonomous regulation. This was found to be true also for *mir-71* expression, with activation of neuronal KGB-1 potently repressing intestinal *mir-71* expression, while activation of the intestinal KGB-1 demonstrated marginal effects, similar to those observed in *kgb-1* mutants (Fig. S1).

KGB-1 activation in larvae also affected *mir-71* expression. However, in agreement with its age-dependent antagonistic contributions, larval activation of KGB-1, contrasting with its activation in adults, increased *mir-71* expression (Fig. 3A and B). This was supported by qRT-PCR measurements demonstrating increased expression of the endogenous non-processed pri-mir-71 (Fig. 3C). In conclusion, *mir-71* expression reflects the age-dependent antagonistic contributions of KGB-1 - induced by KGB-1 activation in larvae but repressed by KGB-1 activation in adults.

**Figure 3.**
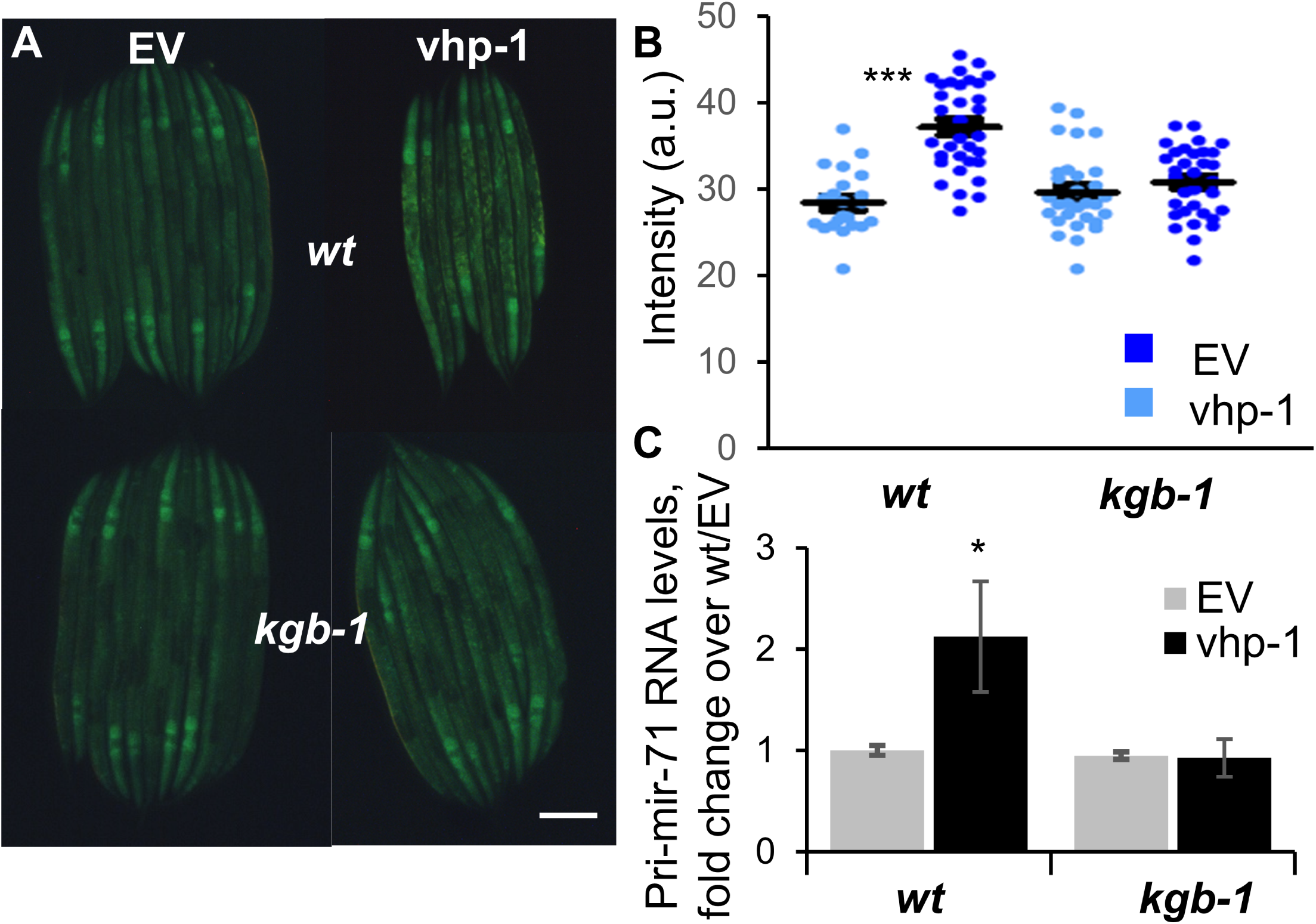
KGB-1 activation in larvae induces mir-71 expression. **(A)** Representative images of transgenic mir-71p::gfp animals and mir-71p::gfp;*kgb-1(km21)* mutants raised from the egg stage to L3 on control (EV) or vhp-1 RNAi. Scale bar, 100 mm **(B)** Quantification of signal intensity in worms as in A. Lines mark averages ± SE; individual measurements shown in dots, 22-33 worms per group; *** p<0.001, t-test. **(C)** qRT-PCR measurements of *mir-71* expression (unprocessed pri-mir-71) in strains and RNAi treatments as designated. Shown are averages and SDs for two independent experiments; * p<0.05.

### mir-71 is required for KGB-1 dependent attenuation of DAF-16 nuclear localization in adults but not for promoting it in larvae

KGB-1 activation was previously shown to modulate DAF-16 nuclear localization in an age-dependent and antagonistic manner, and *daf-16* was shown to be required for most of the KGB-1-dependent phenotypes (Twumasi-Boateng 2012). mir-71 was also shown to modulate DAF-16 nuclear localization (Boulias, 2013). Thus, we examined whether mir-71 was involved in modulating DAF-16 nuclear localization following KGB-1 activation, specifically DAF-16 attenuation following KGB-1 activation in adults. To this end we used transgenic strains expressing a DAF-16::GFP fusion protein from the *daf-16* promoter (Henderson and Johnson 2001). Sterilization of such worms following exposure to cdc-25.1 RNAi led to prominent nuclear localization of DAF-16 (Fig. 4A). This nuclear localization was significantly attenuated following vhp-1 knock-down in wildtype, but not in *kgb-1* mutants, indicating a role for KGB-1 (Fig. 4B). A thwarted effect is also observed in *mir-71* mutants, but this is more difficult to interpret since these mutants show little nuclear localization to begin with. However, qRT-PCR measurements of *mtl-1* expression, a surrogate for DAF-16’s output, showed loss of KGB-1-dependent repression, both in *mir-71*, as well as in *alg-1* mutants, supporting a role for mir-71 in mediating the detrimental effects of KGB-1 activation on DAF-16 output (Fig. 4C). These results suggest that mir-71 mediates KGB-1-dependent DAF-16 regulation in adults and places it upstream of DAF-16.

**Figure 4.**
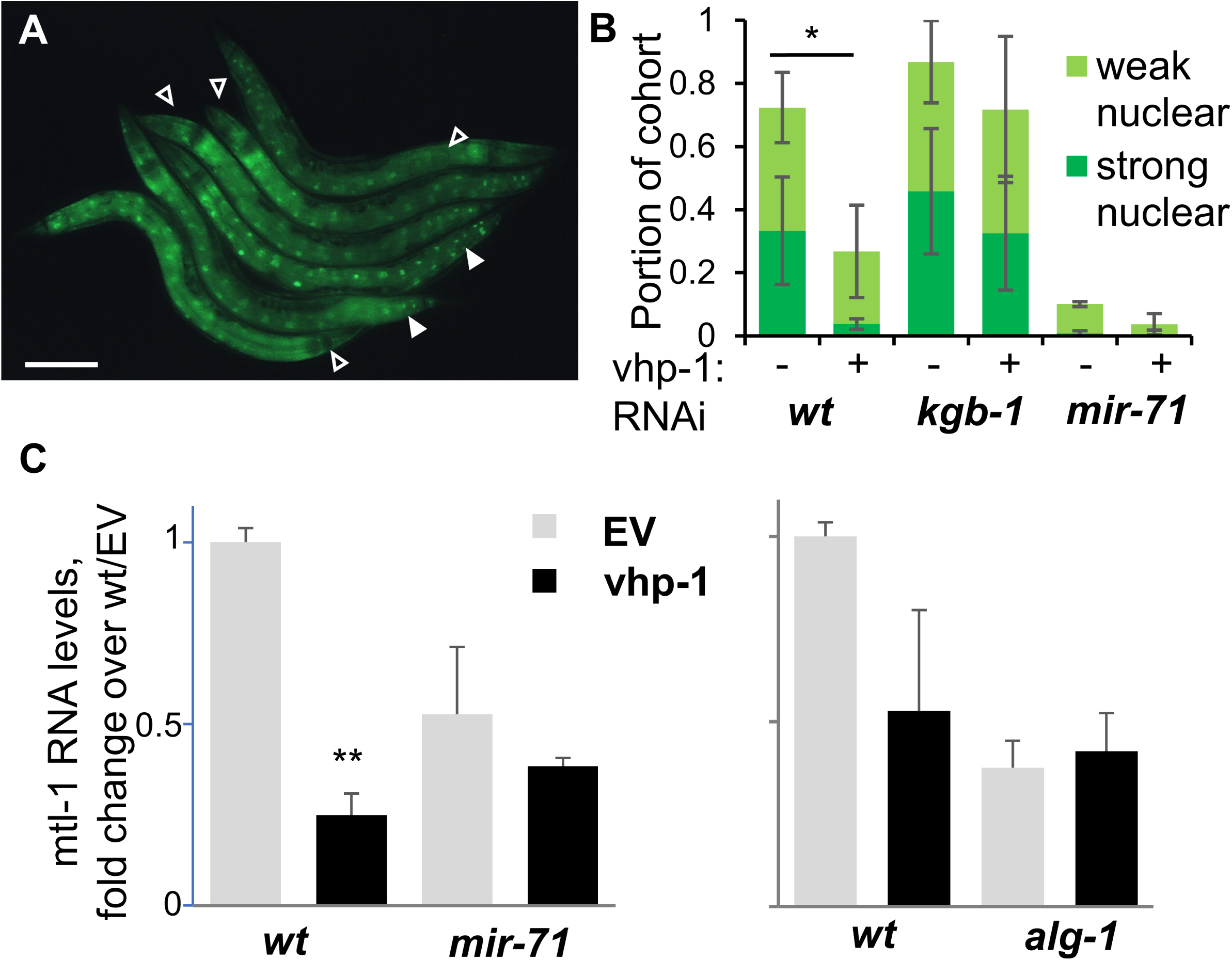
mir-71 mediates effects of KGB-1 activation on DAF-16 output. **(A)** A **r**epresentative image of cdc-25.1(RNAi)-sterilized transgenics expressing a DAF-16::GFP fusion protein following a two-day exposure, beginning at L4, to control RNAi (EV) RNAi. Filled arrowheads mark worms with strong nuclear localization, empty arrowheads mark weak nuclear localization. Scale bar, 200 mm **(B)** Portion of worms with DAF-16::GFP nuclear localization as in A, with genetic backgrounds as designated, and two-day exposure to the designated RNAi. Shown are averages ± SDs of 3 independent experiments (N=346-718 worms total per group); * p<0.05, t-test. **(C)** qRT-PCR measurements of *mtl-1* gene expression in worms of the designated strains, following a two-day exposure to the designated RNAi’s. Shown for each graph are averages ± SDs for 2 independent experiments; **, p<0.01.

In contrast to the role of mir-71 in KGB-1-dependent DAF-16 regulation in adults, mir-71 appeared to be dispensable for enhanced nuclear localization of DAF-16 following KGB-1 activation in larvae (Fig. S2). In agreement, no changes were observed in the induced expression of the DAF-16 target gene *mtl-1* following KGB-1 activation in larvae. Thus, while mir-71 was inversely regulated by KGB-1 in larvae and adults, its contribution to mediating KGB-1 effects on DAF-16 activity was specific to the adult stage, whereas no contribution was observed in larvae.

### MicroRNA processing and mir-71 are required for a subset of KGB-1’s protective contributions in developing larvae

In developing larvae KGB-1 activation was shown to protect animals from protein folding stress in the endoplasmic reticulum (ER stress) and against heavy metals. Protection from heavy metals was shown to be dependent on DAF-16 (Zhang *et al.* 2017), which was also suggested to provide protection from ER stress (Henis-Korenblit *et al.* 2010; Safra *et al.* 2014). To examine the involvement of microRNA processing and mir-71 in these protective contributions, we evaluated development of *alg-1* and *mir-71* mutants in the presence of tunicamycin, an inhibitor of protein glycosylation, which causes ER stress, and their resistance to acute heavy metal stress in the form of 10 mM cadmium. We found that *alg-1* mutants were somewhat susceptible to ER stress, showing retarded development similar to that observed for *daf-16* mutants, although not as dramatic as in *kgb-1* mutants (Fig. 5A). *mir-71* mutants on the other hand were as resistant to ER stress as wildtype animals. This suggested that the microRNA processing machinery was involved in developmental ER stress resistance, but could not account for the full scope of KGB-1’s contribution; further, *mir-71* was dispensable. In contrast to the partial involvement in developmental ER stress resistance, larval resistance to cadmium following KGB-1 activation was fully dependent on *alg-1* and specifically required *mir-71* (Fig. 5B). Together, these results indicate that microRNAs are involved in mediating the protective contributions of KGB-1 in larvae, and that *mir-71* is specifically required for some of these contributions, but redundant for others.

**Figure 5.**
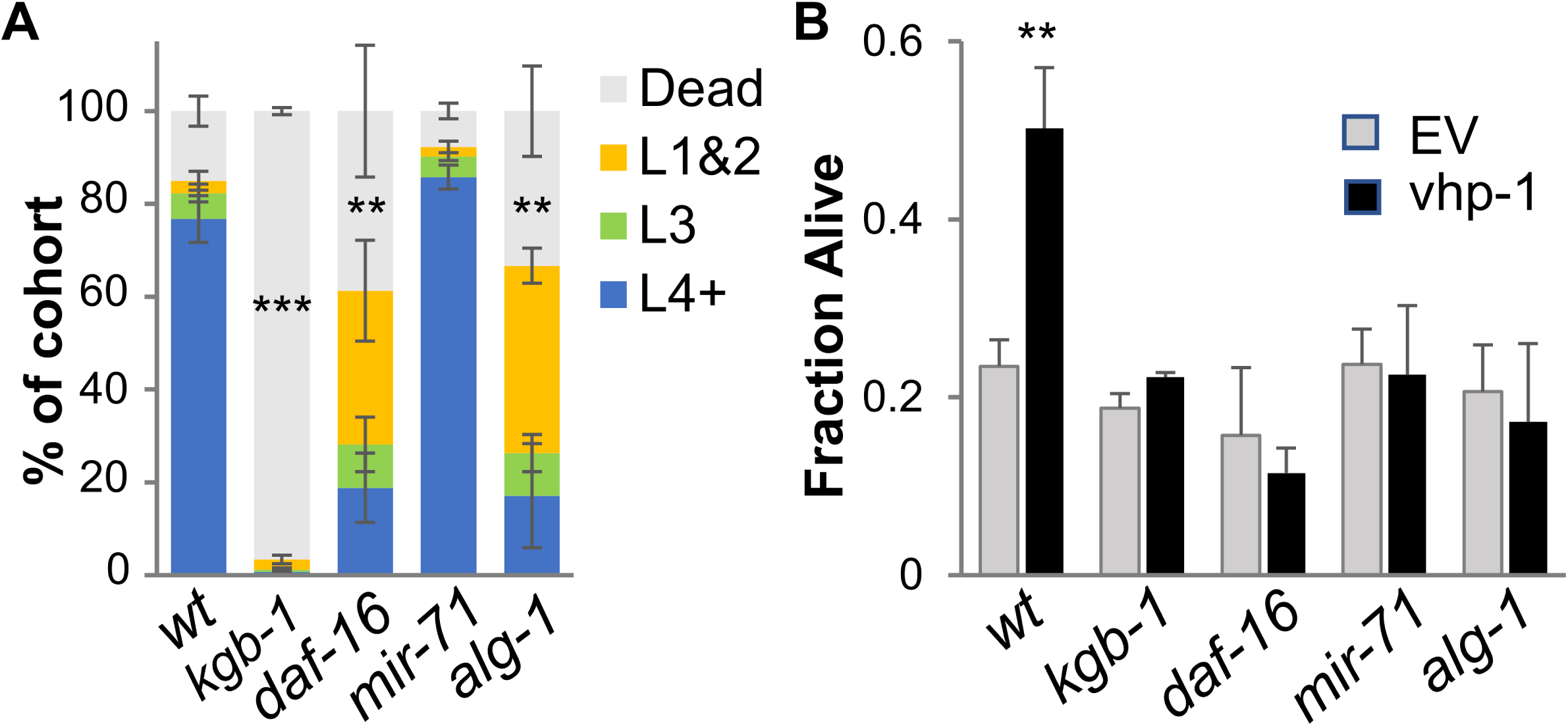
mir-71 is required for some but not all of KGB-1’s protective effects in larvae. **(A)** Development in the presence of 1 mg/ml tunicamycin (3 days, 20°C). Shown are averages ± SDs of two experiments, each performed in duplicate; cohorts included 127-521 worms per replicate. **, p<0.01, ***, p<0.001 (paired t-test), calculated for L4+ worms, with similar results for dead worms. **(B)** Resistance to acute cadmium exposure (10 mM, L3 larvae, 11 hours). Averages ± SDs of two experiments (except for daf-16, which was measured in one experiment only), **, p<0.01 (paired t-test).

## DISCUSSION

Previous work identified an age-dependent switch in the contributions of KGB-1 to stress resistance, which was associated with age-dependent and opposing effects on DAF-16. However, what causes this switch and how KGB-1 activity affected DAF-16 remained a mystery. The results described here offer a clue by identifying the microRNA mir-71 as a downstream mediator of KGB-1, showing age-dependent regulation by KGB-1 and linking KGB-1’s activity both to previously-described phenotypes as well as to DAF-16 output.

The importance of mir-71 as a downstream mediator of KGB-1 is not similar in different ages. Whereas it was fully required for KGB-1’s detrimental contributions in adult animals, i.e. to increased sensitivity to infection, to shortened lifespan, and to reduced DAF-16-dependent expression, it was partially redundant for the beneficial contributions of KGB-1 in developing larvae, i.e. it was required for larval resistance to acute cadmium stress following KGB-1 activation, but not for KGB-1-dependent DAF-16 nuclear localization in larvae and for protection from tunicamycin-induced ER stress. While mir-71 was not required for ER-stress resistance in larvae, disruption of the argonaute gene *alg-1*, involved in microRNA processing and binding, did compromise ER stress resistance, suggesting that other microRNAs beside mir-71 may be involved. Redundancy in microRNA contributions in developing worms is well-described, thought to ensure developmental robustness (Miska *et al.* 2007; Weaver and Han 2018). The partial functional redundancy observed for larval *mir-71* is thus in line with these previous results. However, the observation that mir-71 disruption was sufficient to abolish KGB-1-dependent resistance to acute cadmium exposure may suggest a more central role for mir-71 than other microRNAs in mediating KGB-1-dependent regulation of stress resistance, perhaps through coordination of global miRNA expression (Inukai *et al*. 2018)

Previous studies have shown that mir-71 extended lifespan, primarily through repression of insulin signaling (de Lencastre *et al.* 2010; Zhang *et al.* 2011). In agreement with this, mir-71 was subsequently shown to promote nuclear localization and activity of DAF-16 in the intestine following germ cell loss, only that this contribution depended on neuronal expression of mir-71 (Boulias and Horvitz 2012). Support for neuronal functions of mir-71 was further provided by studies showing both neuronal cell autonomous repression of TIR-1/SARM - an upstream regulator of the p38 pathway, as well as cell non-autonomous contributions of neuronal mir-71 and TIR-1 to intestinal proteostasis (Couillault *et al.* 2004; Hsieh *et al.* 2012; Finger *et al.* 2019). It is not clear whether the DAF-16-dependent lifespan extension and TIR-1-dependent proteostasis are related, but to date no evidence exists for interactions between the two important regulators. Our results support a role for mir-71 in regulation of DAF-16 and suggest that this regulation can be modulated by KGB-1. Whereas our results do not resolve the tissue from which mir-71 exerted its effects, they do show that it is expressed in the intestine, and that this expression is regulated by KGB-1 in accordance with KGB-1’s age-dependent contributions. Whether this intestinal regulation is downstream to the effects of mir-71 on DAF-16 and on survival phenotypes (as some sort of a feedback loop), or upstream of these effects, suggesting a causative role, is not known. Interestingly, our results suggest that mir-71 is involved in cell nonautonomous regulation, but in a different way than those suggested before, as intestinal mir-71 expression depends primarily on neuronal KGB-1, much like many of the KGB-1-dependent phenotypes (Fig. S1)(Liu *et al.* 2018).

It is worth noting that additional targets were described to be repressed by mir-71, including the transcription factor PHA-4, and the argonaute ALG-1 (Smith-Vikos *et al.* 2014; Inukai *et al.* 2018). The antagonistic nature of these interactions predicts that disruption of *mir-71* and *alg-1* will have opposing effects on downstream phenotypes. However, the results described here demonstrate similar effects of *mir-71* and *alg-1* disruptions on lifespan, DAF-16 nuclear localization and on cadmium resistance, indicating that ALG-1 repression is likely not playing a role in the KGB-1-dependent regulation and phenotypes described here.

The work here advances our understanding of the antagonistic pleiotropy behavior of KGB-1 one notch forward. It points at mir-71 as a hub for age-dependent regulation – both of DAF-16, and potentially of additional targets. It further demonstrates age-dependent regulation of mir-71 by KGB-1, but the mechanism responsible for this remains to be found.

## ACKNOWLEDGEMENTS

We thank the *Caenorhabditis* Genetics Center and Dr. Zachary Pincus from Washington University, St. Louis for strains. Work on this project was supported by an NSF grant, IOS-1355240.

## AUTHOR CONTRIBUTIONS

C.R. designed and performed experiments, carried out analysis, and summarized results. M.S. initiated the project, designed and performed experiments and summarized results. Both took part in the writing of the manuscript.

## FIGURE LEGENDS

**Figure S1.**
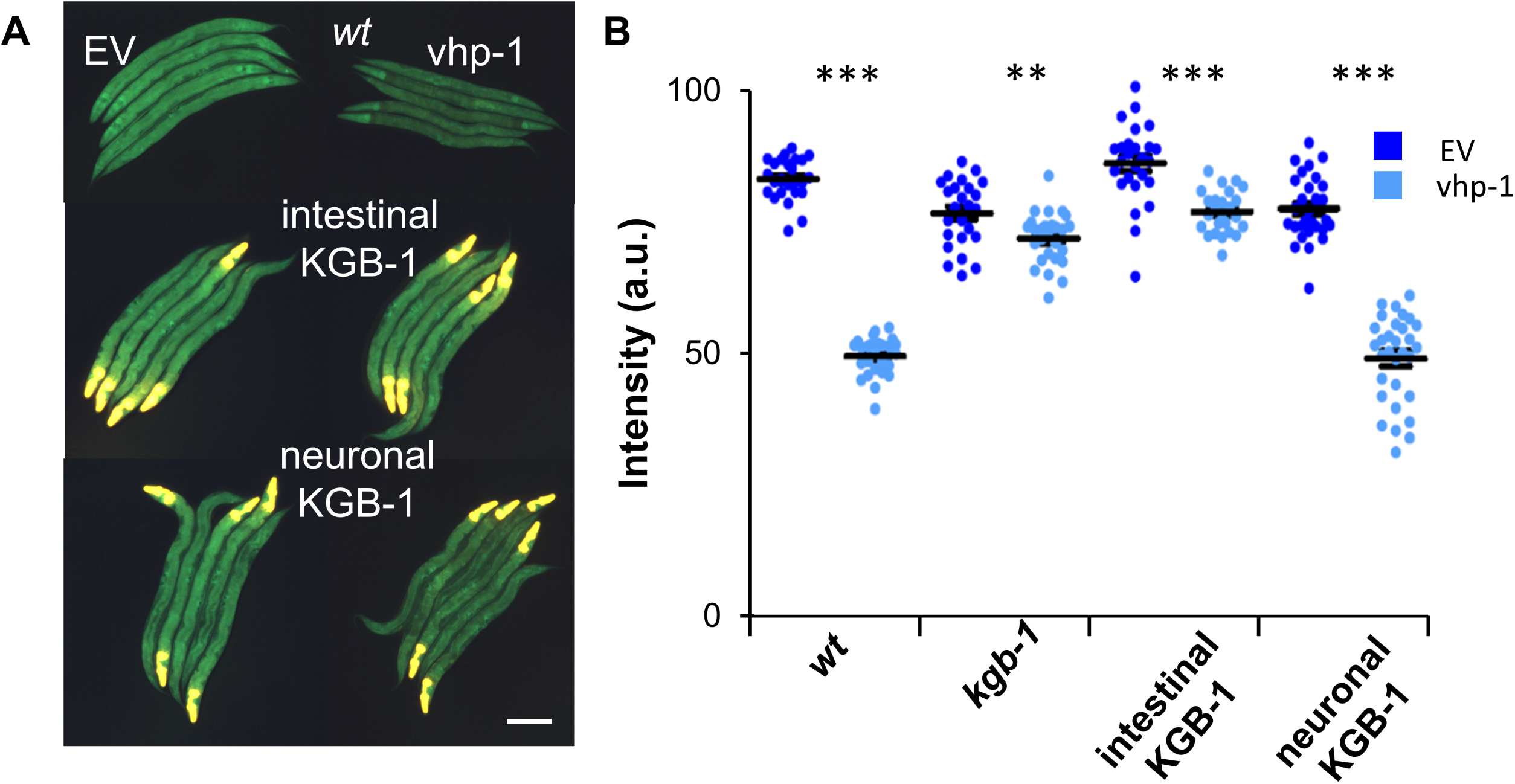
KGB-1 suppresses mir-71 expression cell non-autonomously. **(A)** Representative images of cdc-25.1(RNAi)-sterilized transgenic animals expressing GFP from the *mir-71* promoter and kgb-1 from tissue-specific promoters, as designated, following a 2-day exposure, beginning at L4, to control (EV) or vhp-1 RNAi. Pharyngeal yellow fluorescence results from bleed through of a myo-2p::tdTomato co-injection marker into the green channel – this area was avoided in signal quantification. Scale bar, 200 mm (B) Quantification of signal intensity in worms such as in A. Lines mark averages ± SE; individual measurements shown in dots, 25-31 worms per group; ** p<0.01, *** p<0.001, t-test.

**Figure S2.**
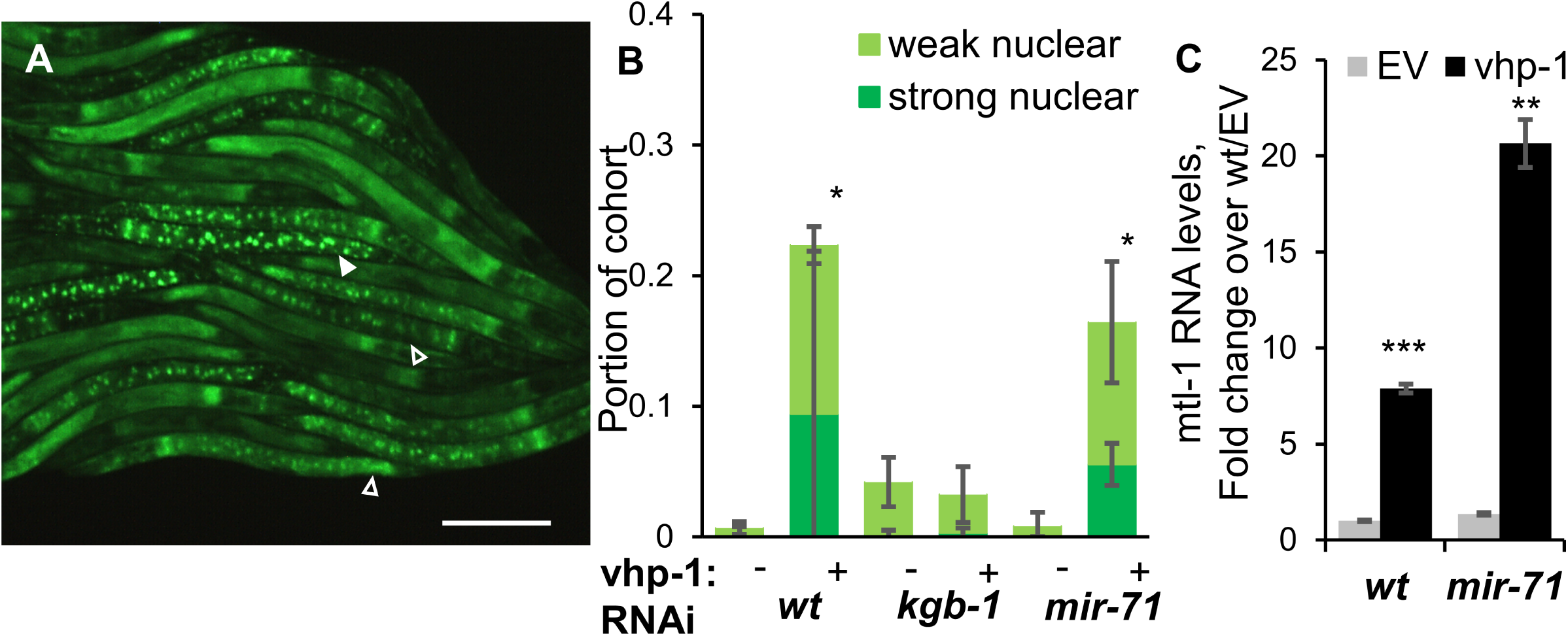
mir-71 is not required for KGB-1 dependent enhancement of DAF-16 function in larvae. **(A)** A representative image of L3 transgenic animals expressing a DAF-16::GFP fusion protein following exposure, from egg stage to control RNAi (EV) RNAi. Filled arrowheads mark worms with strong nuclear localization, empty arrowheads mark weak nuclear localization. Scale bar, 100 mm (B) Portion of worms with DAF-16::GFP nuclear localization as in A, with genetic backgrounds and RNAi treatments as designated. Shown are averages and SDs of 3 independent experiments (N=346-718 worms total per group); * p<0.05, t-test. (C) qRT-PCR measurements of *mtl-1* gene expression in L4 larvae of strains with RNAi treatments as designated. Shown are averages ± SDs for 2 independent experiments.

**Table S1.**
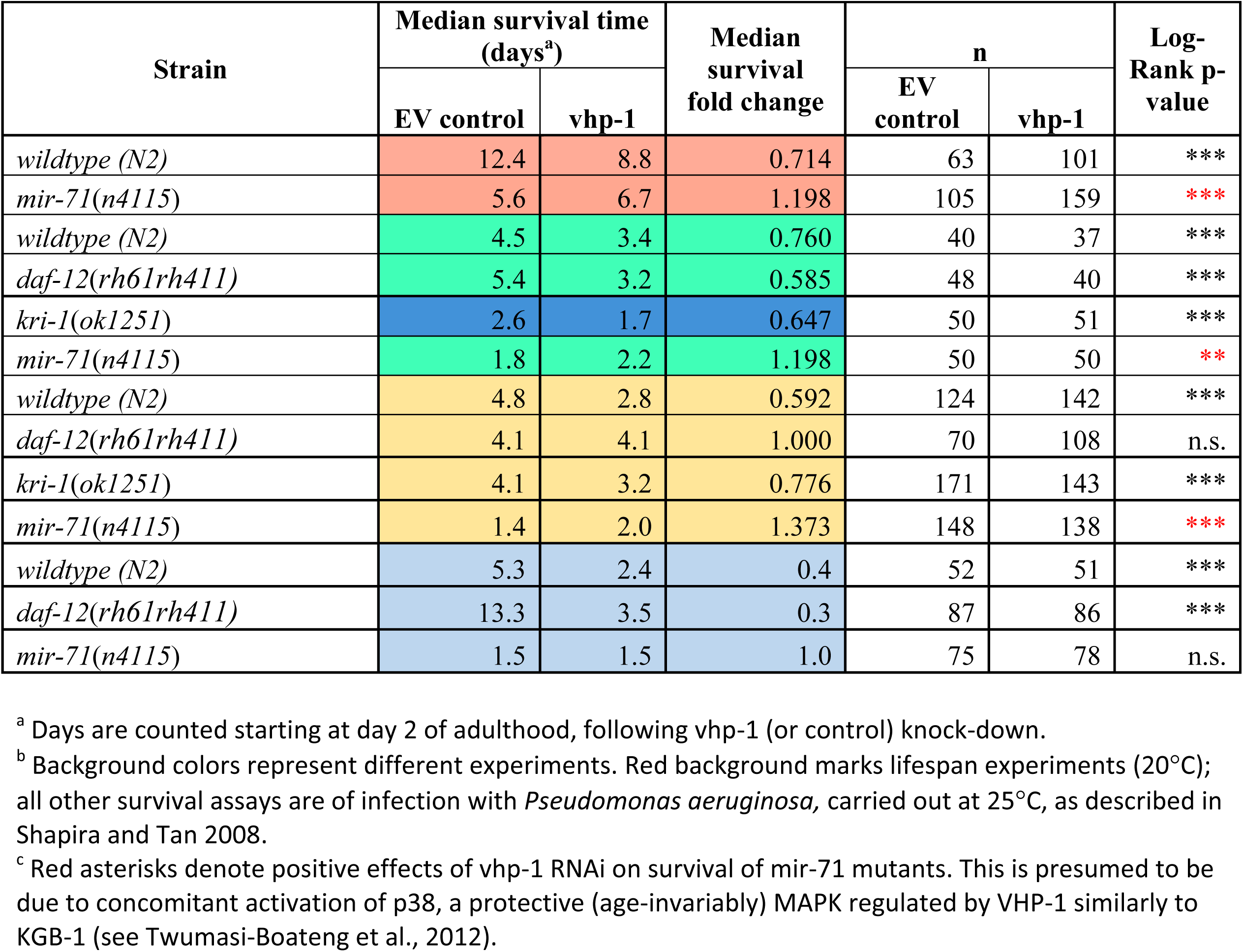
Detrimental effects of KGB-1 depend on mir-71.

| Strain | Median survival time (days<sup>a</sup>) |  | Median survival fold change | n |  | Log-Rank p-value |
| --- | --- | --- | --- | --- | --- | --- |
|  | EV control | vhp-1 |  | EV control | vhp-1 |  |
| *wildtype (N2)* | 12.4 | 8.8 | 0.714 | 63 | 101 | *** |
| *mir-71(n4115)* | 5.6 | 6.7 | 1.198 | 105 | 159 | *** |
| *wildtype (N2)* | 4.5 | 3.4 | 0.760 | 40 | 37 | *** |
| *daf-12(rh61rh411)* | 5.4 | 3.2 | 0.585 | 48 | 40 | *** |
| *kri-1(ok1251)* | 2.6 | 1.7 | 0.647 | 50 | 51 | *** |
| *mir-71(n4115)* | 1.8 | 2.2 | 1.198 | 50 | 50 | ** |
| *wildtype (N2)* | 4.8 | 2.8 | 0.592 | 124 | 142 | *** |
| *daf-12(rh61rh411)* | 4.1 | 4.1 | 1.000 | 70 | 108 | n.s. |
| *kri-1(ok1251)* | 4.1 | 3.2 | 0.776 | 171 | 143 | *** |
| *mir-71(n4115)* | 1.4 | 2.0 | 1.373 | 148 | 138 | *** |
| *wildtype (N2)* | 5.3 | 2.4 | 0.4 | 52 | 51 | *** |
| *daf-12(rh61rh411)* | 13.3 | 3.5 | 0.3 | 87 | 86 | *** |
| *mir-71(n4115)* | 1.5 | 1.5 | 1.0 | 75 | 78 | n.s. |

<sup>a</sup> Days are counted starting at day 2 of adulthood, following vhp-1 (or control) knock-down.
<sup>b</sup> Background colors represent different experiments. Red background marks lifespan experiments (20°C); all other survival assays are of infection with *Pseudomonas aeruginosa*, carried out at 25°C, as described in Shapira and Tan 2008.
<sup>c</sup> Red asterisks denote positive effects of vhp-1 RNAi on survival of mir-71 mutants. This is presumed to be due to concomitant activation of p38, a protective (age-invariably) MAPK regulated by VHP-1 similarly to KGB-1 (see Twumasi-Boateng et al., 2012).

**Table S2.**
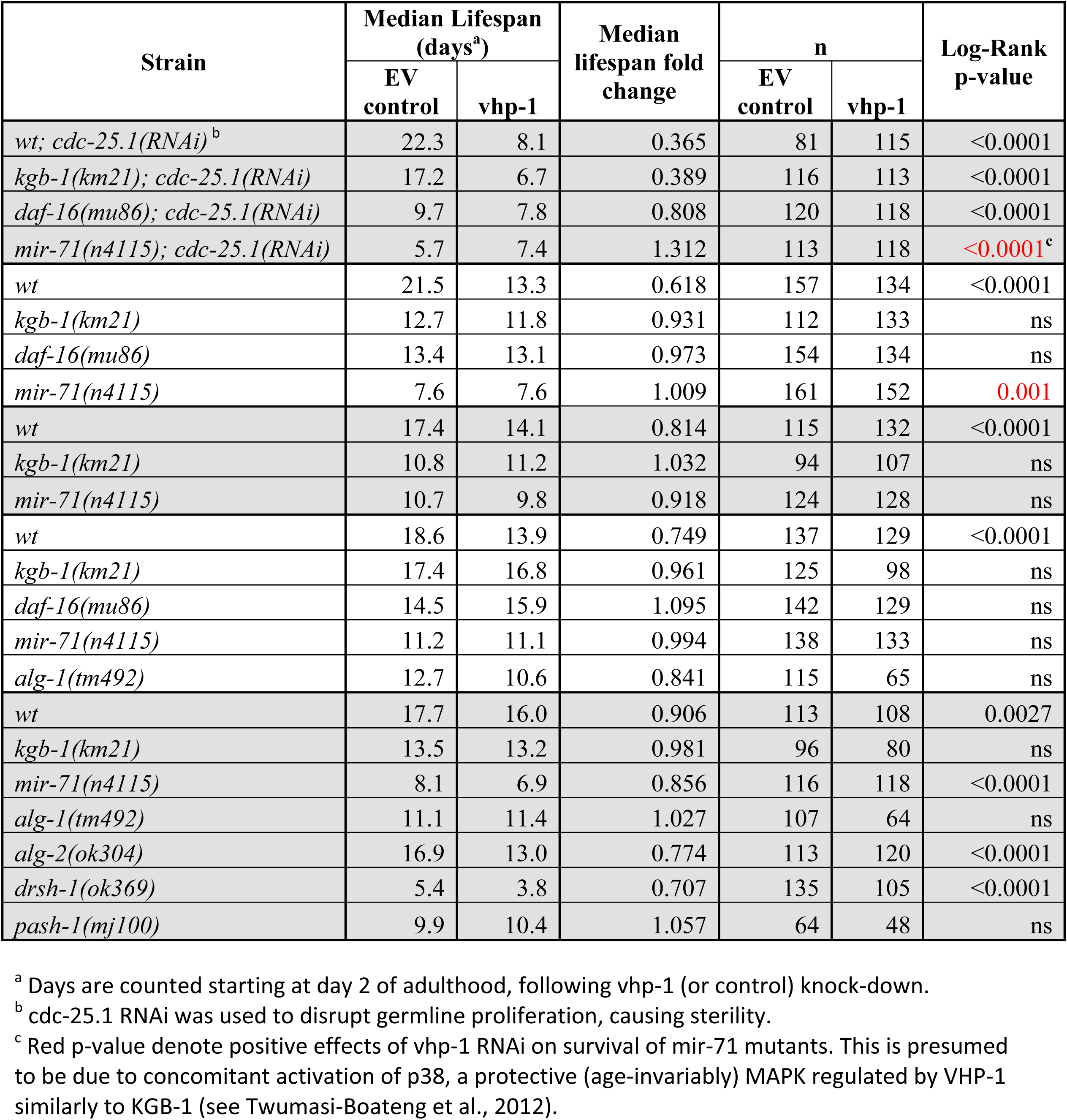
Involvement of *mir-71* and the microRNA machinery in KGB-1 effects in fertile and sterile worms.

